# The energetics of rapid cellular mechanotransduction

**DOI:** 10.1101/2022.07.17.500355

**Authors:** Michael N. Young, Michael J. Sindoni, Amanda H. Lewis, Stefan Zauscher, Jörg Grandl

## Abstract

Cells throughout the human body detect mechanical forces. While it is known that the rapid (millisecond) detection of mechanical forces is mediated by force-gated ion channels, a detailed quantitative understanding of cells as sensors of mechanical energy is still lacking. Here, we combine atomic force microscopy with patch-clamp electrophysiology to determine the physical limits of cells expressing the force-gated ion channels Piezo1, Piezo2, TREK1, and TRAAK. We find that, depending on the ion channel expressed, cells can function either as proportional or nonlinear transducers of mechanical energy, detect mechanical energies as little as ∼100 fJ, and with a resolution of up to ∼1 fJ. These specific energetic values depend on cell size, channel density, and cytoskeletal architecture. We also make the surprising discovery that cells can transduce forces either nearly instantaneously (< 1ms), or with substantial time delay (∼10 ms). Using a chimeric experimental approach and simulations we show how such delays can emerge from channel-intrinsic properties and the slow diffusion of tension in the membrane. Overall, our experiments reveal the capabilities and limits of cellular mechanosensing and provide insights into molecular mechanisms that different cell types may employ to specialize for their distinct physiological roles.

## Introduction

The ability to detect mechanical forces is essential for a breadth of physiological processes including our sense of light touch, proprioception, blood pressure regulation, bone homeostasis, interoception, and cell differentiation (Li et al., 2019; Pathak et al., 2014; Ranade et al., 2014b; Sun et al., 2019; Woo et al., 2014, 2015; Zeng et al., 2018). Despite this broad importance, our quantitative understanding of cells as sensors of mechanical energy is extremely limited and even seemingly simple questions about cellular mechanotransduction remain unanswered: What is the smallest mechanical energy a cell can detect? What is the smallest mechanical energy a cell can resolve? And how fast can cells respond?

The most rapid detection of forces is mediated by force-gated ion channels (FGICs), which respond in as little as 40 µs by gating the flux of ions (Corey and Hudspeth, 1979). FGICs vary in their ion selectivity, single-channel conductance, and gating kinetics, which enables cells to transduce mechanical forces into distinct electrochemical signals. However, how FGICs compare in their most fundamental property, the sensing of mechanical energy, is not well understood (Young et al., 2022). For example, measurements of the gating energy of the well-studied ion channel Piezo1 differ substantially (8·10^−21^ J and 40·10^−21^ J), while for the two-pore potassium channels TREK1 and TRAAK estimates span 1-10·10^−21^ J, and for Piezo2 and the recently discovered TMEM63 channels such information is missing altogether (Brohawn et al., 2014a; Cox et al., 2016; Lewis and Grandl, 2015; Maksaev et al., 2011; Murthy et al., 2018).

Similarly, fundamental molecular mechanisms of many FGICs are still unclear: while some FGICs, such as Piezo1, TREK1, TREK2, and TRAAK, can be directly activated by membrane tension, others such as NOMPC sense force through tethers (Aryal et al., 2017; Berrier et al., 2013; Brohawn et al., 2014b; Cox et al., 2016; Lewis and Grandl, 2015; Maingret et al., 1999; Syeda et al., 2016; Zhang et al., 2015). Importantly, both mechanisms are not mutually exclusive and may act in synergy (Sukharev and Sachs, 2012).

Structural and mechanical properties of the cell are tightly coupled to force detection. Specifically, the actin cytoskeleton can absorb mechanical forces (mechanoprotection), while also transmitting forces near or directly to the FGIC (mechanotransmission). For example, the actin cytoskeleton is a major determinant of cell stiffness and slows the transmission of membrane tension (Grady et al., 2016; Shi et al., 2018). In contrast, the actin cytoskeleton can tether to Piezo1 via the cadherin-ß-catenin complex so that actin disruption decreases Piezo1-mediated responses to cell indentation. Additionally, the actin cytoskeleton has been proposed to transmit long-range forces to Piezo2 channels (Gottlieb et al., 2012; Verkest et al., 2022; Wang et al., 2022). Effectively, both processes compete and therefore to what extent the actin cytoskeleton facilitates or impedes force sensing remains an open question.

Another impediment to a quantitative understanding of cells as mechanotransducers are technical limitations: The standard assays for probing cellular mechanotransduction use stimulation by pressure-clamp (stretch) and cell-indentation (poke) in combination with patch-clamp electrophysiology (Besch et al., 2002; Hao and Delmas, 2011). While both methods provide an accurate readout of the electrical response, stretch stimulation disrupts the underlying cytoskeleton and does not inform about the overall cellular response, and poke stimulation cannot quantify the magnitude of the applied force. More sophisticated methods for mechanical stimulation include magnetic force actuators, atomic force microscopy, optical tweezers, and ultrasound, but often are combined with calcium-imaging, which does not yield a precise measure of channel activity or kinetics and precludes the study of FGICs that do not conduct calcium (Falleroni et al., 2018; Gaub and Müller, 2017; Hoffman et al., 2022; Jembrek et al., 2015; Lee et al., 2014; Lin et al., 2019; Poole et al., 2014; Prieto et al., 2018; Ranade et al., 2014a; Sorum et al., 2021; Wu et al., 2016; Ye et al., 2018). In summary, none of these assays can stimulate cells with a precisely quantified indentation, force, and energy, while also measuring the transduction current with the gold-standard of electrophysiology.

Here, we developed an instrument that combines atomic force microscopy (AFM) with patch-clamp electrophysiology to allow for quantitative mechanical stimulation and simultaneous detection of the evoked transduction current. With this tool we set out to explore and precisely quantify how single cells convert the energy of mechanical compression into an electric signal.

## Results

### Simultaneous measurements of cell compression and mechanotransduction

To quantify the magnitude of energies that cells can sense and respond to we built an instrument that combines an atomic force microscope (AFM) with patch-clamp electrophysiology (**Figure 1A-C**). In each experiment, a flexible cantilever compresses a single cultured cell at a speed of 40 µm/s, and then holds its position for 100 ms, before it is retracted at the same speed. The cantilever, whose dimensions are comparable to the size of the cell, is centered on a cell, and displaced for a total of ∼6 µm, which altogether results in a large-scale compression of the entire cell. Since the stiffness of each cantilever and the sensitivity of the detector to cantilever bending is calibrated for individual experiments, we can calculate the distance (*d*), compression force (*F*), and cumulative work (*W*) performed on the cell throughout the entire experiment with a relative uncertainty of ≲ 20% (see Methods). At the same time, we record electrophysiologically from the cell by patching it in the whole-cell configuration, which enables us to measure the evoked transduction current (*I*) (**Figure 1D**).

**Figure 1:**
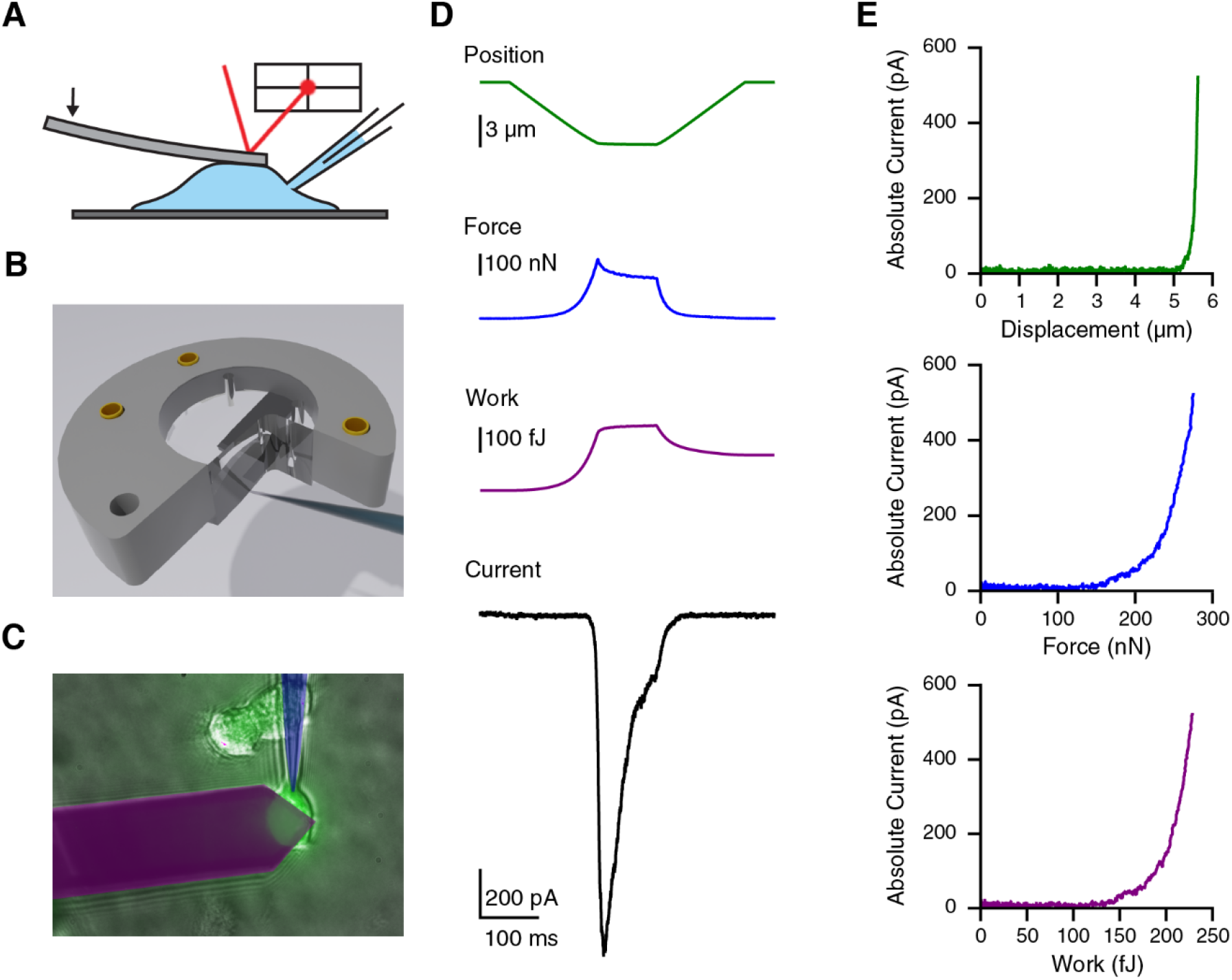
A new instrument to simultaneously measure mechanical work and transduction. **(A)** Schematic of AFM-Electrophysiology setup. **(B)** 3D schematic showing modified probe holder allowing access to the cell by a pipette. **(C)** Image of cantilever and pipette over a YFP-labeled cell as seen in a typical experiment. **(D)** Representative trace of a Piezo1-expressing cell in response to a typical AFM-electrophysiology protocol. **(E)** Absolute current as a function of distance (*top; green*), force (*middle; blue*), work (*bottom; purple*).

Initial validation experiments on HEK293T cells overexpressing the force-gated ion channel Piezo1 show that AFM compression evokes rapid transduction currents at -80 mV, which are absent in cells transfected with YFP as control. Peak current amplitudes (1.1±0.5 nA, n=16) are comparable to those from classical poke-indentation experiments and inactivate with a time-course of 25±4 ms (n=16), which is expected for Piezo1 at this holding potential (Coste et al., 2010; J. Wu et al., 2017). We observed that the cantilever deflection is undergoing a small, but noticeable adaptation during the static phase with a time course of 33±3 ms (n=16) when fit with a single exponential decay, perhaps due to relaxation of the cell. We therefore focused all subsequent analyses on the approach phase to calculate distance-current (*d/I*), force-current (*F/I*), and work-current (*W/I*) relationships for understanding how cell compression evokes mechanotransduction (**Figure 1E**).

### Linearity of cellular mechanotransduction

We next performed systematic AFM-electrophysiology experiments on HEK293T cells overexpressing the force-gated ion channels Piezo1, Piezo2, TREK1, and TRAAK (**Figure 2A, Figure 2—figure supplement 1)**. We first noticed that work-current relationships differed qualitatively in their linearity: cells expressing TREK1 and TRAAK showed currents that were approximately proportional to the work performed on the cell. In comparison, work-current relationships from cells expressing Piezo1 and Piezo2 appeared nonlinear. This qualitative difference suggests that cells may in principle function either as proportional or nonlinear transducers of mechanical energy. For quantification, we fit all individual measurements with a power law (*I* = *W*^*α*^), where a power-coefficient α = 1 indicates a perfectly linear (directly proportional) transduction of mechanical work into current (**Figure 2B**). Consistent with our initial observation, work-current relationships of cells expressing TREK1 and TRAAK have power-coefficients of 1.4±0.1 (n=60) and 1.9±0.2 (n=38), respectively, confirming that cells expressing TREK1 and to a lesser extent TRAAK behave approximately as proportional transducers of mechanical energy. However, cells expressing Piezo2, and even more so, cells expressing Piezo1 have substantially larger power coefficients of 2.6±0.2 (n=47), and 6.2±0.4 (n=39), respectively, meaning that their responses are indeed highly nonlinear, i.e., their force-detection is akin to a switch. More generally, the data demonstrate that the identity of the transduction channel defines this important response property.

**Figure 2.**
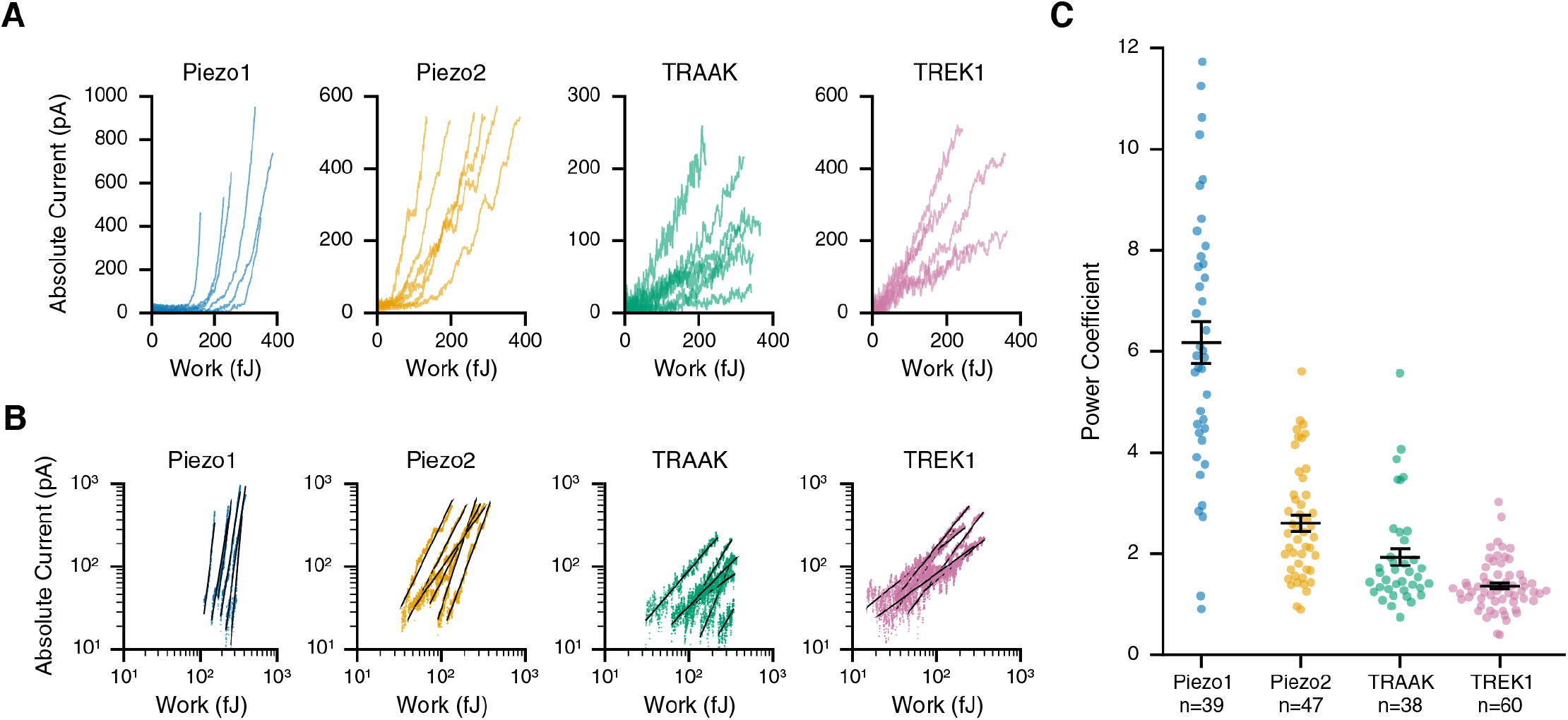
Linearity of cellular mechanotransduction. **(A)** Raw work-current relationships for six representative cells overexpressing Piezo1 (blue), Piezo2 (orange), TRAAK (green), and TREK1 (pink). Currents are represented as absolute values. **(B)** Respective work-current relationships on double-log scaled axes with power law fits (black) for cells in (A). Data are limited to points beyond determined work thresholds (see Methods). **(C)** Power-coefficients of power-law fits. Error bars indicate mean ± sem; n indicates the number of replicates (individual cells).

### The threshold and resolution for detecting mechanical energy

To quantify the transduction response further, we next calculated for each individual cell its detection threshold (*W*_threshold_), or the mechanical energy required to elicit a detectable transduction current (**Figure 3A**). While measurements vary substantially between individual cells, summary data show that each channel type also confers distinct detection thresholds (**Figure 3B**). Cells expressing TREK1 exhibit the lowest thresholds of 68.2±6.3 fJ (n=60), whereas cells expressing Piezo1 exhibit the highest thresholds of 213.7±16.6 fJ (n=39). In addition, the detection thresholds of cells with the respectively related channels TRAAK (124.5±11.9 fJ; n=38) and Piezo2 (86.8±7.1 fJ; n=47) differed by roughly two-fold from their orthologs, meaning that detection thresholds are not conserved within channel families. Notably, in this analysis the current amplitude corresponding to *W*_*threshold*_ is equivalent to the activation of only ∼2 ion channels (see Methods, (Blin et al., 2016; Coste et al., 2015)). Thus, considering that activating two Piezo1 channels requires an energy of ΔG ∼80·10^−8^ fJ (Cox et al., 2016; Lewis and Grandl, 2015) directly implies that >10^7^-times more energy is absorbed by the cell before transduction is initiated. This result shows a considerable capacity of the cell to buffer mechanical energy.

**Figure 3.**
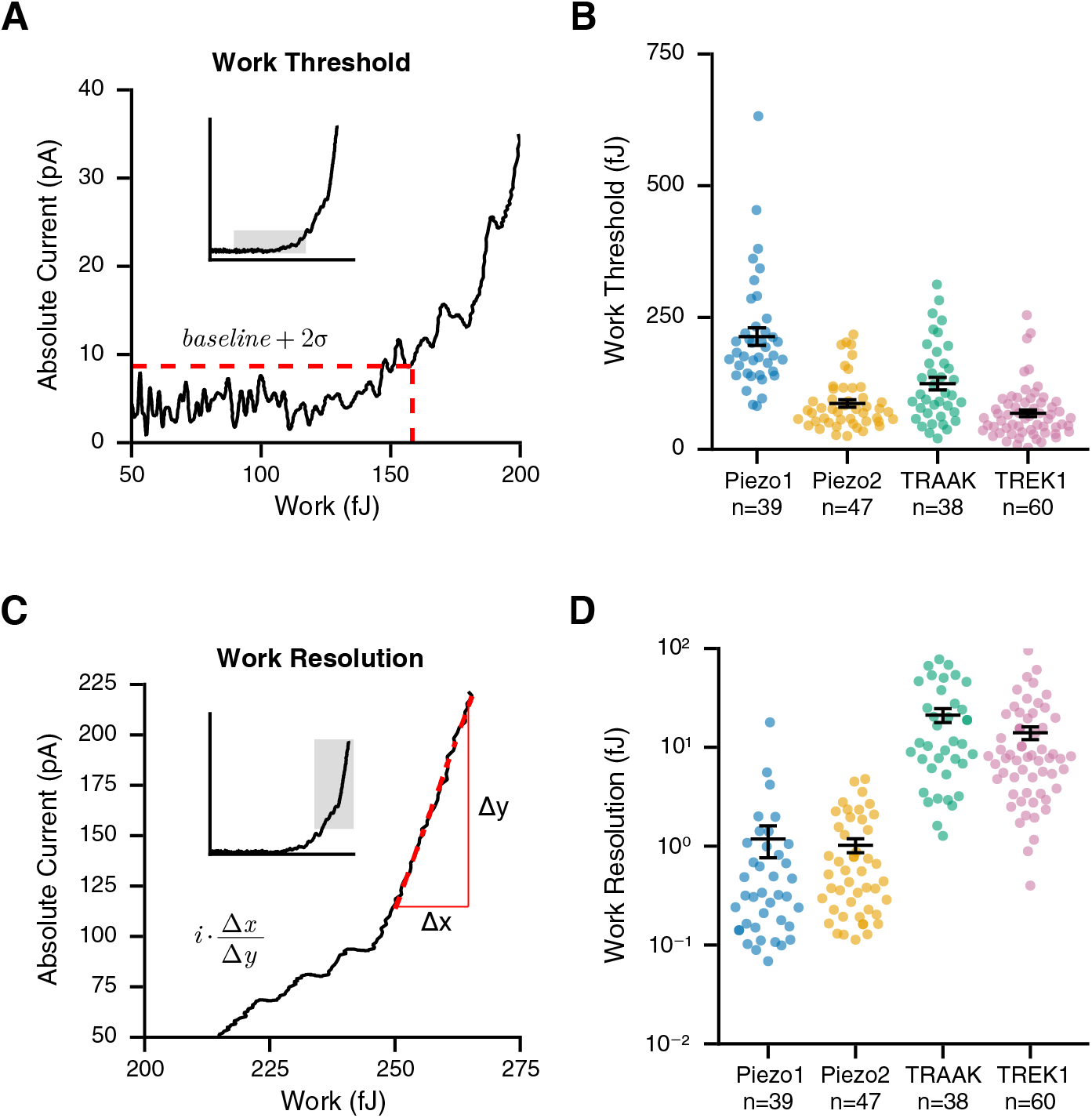
Cellular threshold and resolution for detecting mechanical energy. **(A)** Calculation of work threshold: representative work-current relationship with red, dashed lines indicating the current threshold (average baseline + 2σ) and the corresponding work threshold, respectively. (**B)** Work threshold values for cells overexpressing Piezo1 (blue), Piezo2 (orange), TRAAK (green), and TREK1 (pink). **(C)** Calculation of work resolution: representative work-current relationship with a red, dashed line indicating the maximal slope (Δx/Δy), which is then multiplied by the single-channel current (*i*). **(D)** Work resolution values for cells overexpressing Piezo1 (blue), Piezo2 (orange), TRAAK (green), and TREK1 (pink). Error bars indicate mean ± sem; n indicates the number of replicates (individual cells).

We next reasoned that once this initial buffering capacity is depleted, the subsequent activation of FGICs may be more efficient. To test this idea, we quantified the work resolution (*W*_*resolution*_), which we define as the inverse maximum slope of the work-current relationship normalized by the unitary current of the transduction channel (see Methods, **Figure 3C**). Intuitively, this measure can be understood as the minimal mechanical work required to open the very next ion channel, or as the smallest mechanical energy a cell can discriminate. Indeed, we found that values for *W*_*resolution*_ were almost 100-fold smaller compared to *W*_*threshold*_, and in addition, unlike detection thresholds, similar between related ion channels. Specifically, values for *W*_*resolution*_ are low for cells expressing Piezo1 (1.2±0.4 fJ; n=39) and Piezo2 (1.0±0.2 fJ; n=47), but an order of magnitude higher for cells expressing TRAAK (21.2±3.4 fJ; n=38) and TREK1 (14.0±2.1 fJ; n=60) (**Figure 3D**). Analogous to our above estimate, comparing the gating energy of Piezo1 yields that even when cellular mechanotransduction is at its most sensitive, >10^5^-times more energy is absorbed by the cell than is required for gating the very next ion channel. Altogether, we conclude from this analysis that detection of mechanical energy by cells, despite overexpression of FGICs, is a process that is extremely inefficient.

### Molecular and cellular determinants of work threshold and resolution

Our above measurements of work threshold and work resolution clearly show that the specific values are dependent on the identity of the FGIC and can be quite distinct from each other. However, each specific measurement also exhibits a large degree of variability, which led us to investigate further what other molecular or cellular factors may contribute towards the cellular detection of mechanical energy. First, we reasoned that cell size could play an important role, because it may determine the capacity to buffer mechanical energy. To test this idea, we took advantage of the natural variability in cell size and correlated values for *W*_*threshold*_ and *W*_*resolution*_ of all individual cells with their respective electric capacitance, which scales with membrane area and is thus a proxy for cell size (Hille, 2001). Indeed, for both Piezo1 and Piezo2 values for *W*_*threshold*_ are positively correlated with cell capacitance, supporting this idea (**Figure 4A**).

**Figure 4.**
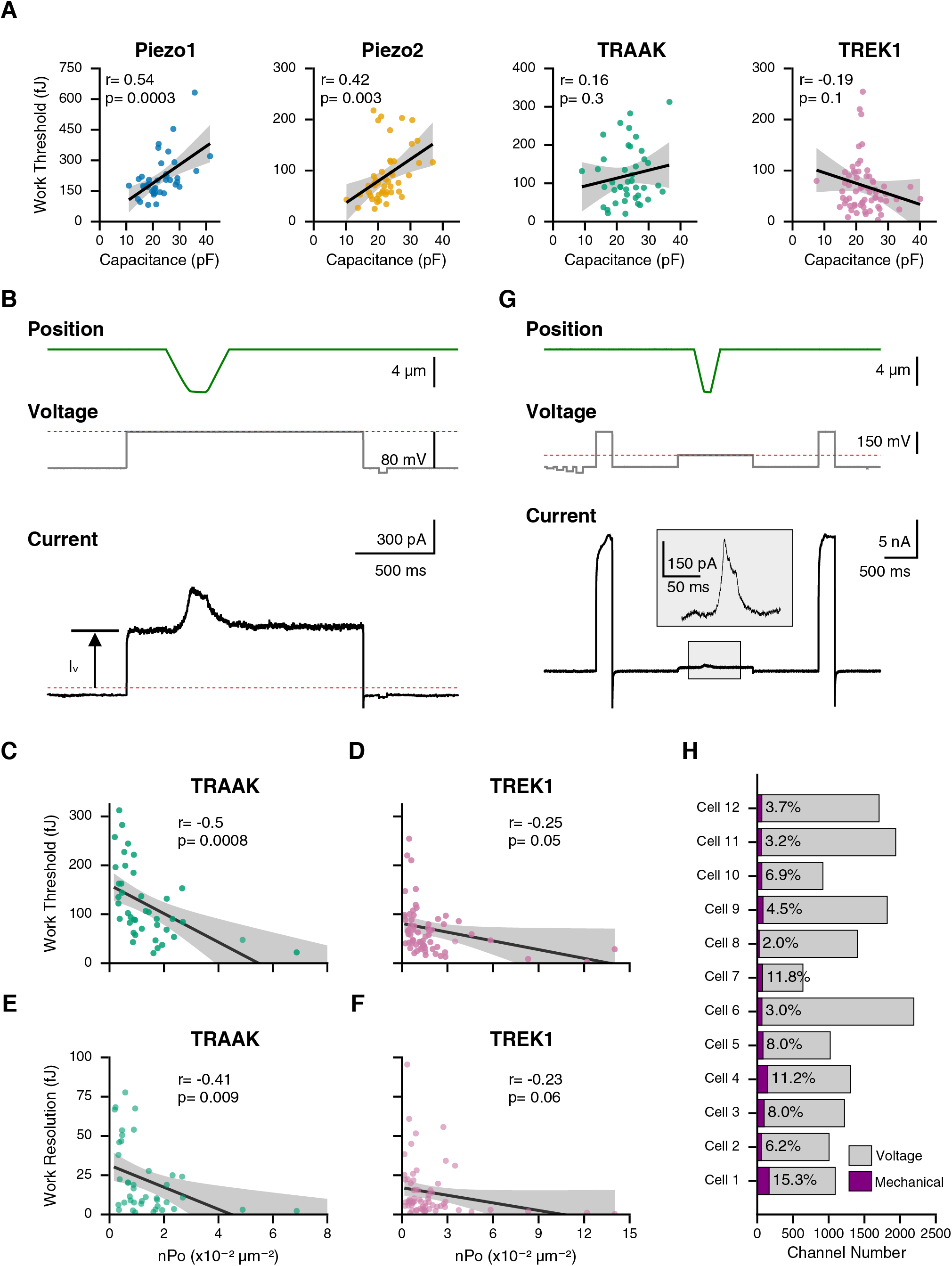
Channel density modulates the cellular threshold and resolution. **(A)** Correlation between values for work threshold and cell capacitance for Piezo1 (blue), Piezo2 (orange), TRAAK (green), TREK1 (pink). Black lines are linear fits and gray bands 95% confidence intervals. Correlation coefficients and p-values are indicated on the top left. **(B)** Protocol to measure mechanotransduction and channel density, and representative current trace. The absolute current at 0 mV (I_V_) was used to determine channel number (N). The red, dashed line indicates zero current. Work threshold and work resolution values as a function of channel density (n·P_o_) for cells overexpressing TRAAK **(C and E)** and TREK1 **(D and F)**, respectively. Black lines are **l**inear fits and gray bands 95% confidence intervals. Correlation coefficients and p-values are indicated on the top right. **(G)** Protocol to measure mechanotransduction and TREK1 channel number, and representative current trace. The inset shows an enlarged view of the mechanotransduction current. Voltage steps to +160 mV were used to count total channel number and mechanocurrent was measured at 0 mV. **(H)** Channel number activated by voltage (gray) and mechanical stimulation (purple) for twelve individual cells. The percentage of TREK1 channels activated by the mechanical stimulus is shown in each bar.

Surprisingly, K2P channel thresholds as well as *W*_*resolution*_ across all channels did not correlate with cell capacitance (data not shown), indicating that cell size is important, but that additional factors may contribute as well. We therefore turned our attention to channel density, which we reasoned might be a more direct predictor of work threshold and work resolution, because it determines the probability of activating multiple channels simultaneously. To test this idea, we performed an additional analysis taking advantage of the polymodal nature of TREK1 and TRAAK, which can be activated by both mechanical force and voltage (Berrier et al., 2013; Maingret et al., 1999). Specifically, we measured the membrane capacitance as a proxy for total membrane surface area and the baseline current at 0 mV to calculate channel density, in addition to probing mechanotransduction as described above (**Figure 4B**, see Methods). In these measurements, channel density varied ∼20-fold for TRAAK and ∼50-fold for TREK1, allowing us to sample a wide range. We indeed found that values for *W*_*threshold*_ and *W*_*resolution*_ correlate negatively with channel density (**Figure 4C-F**). In summary, we conclude that absolute cell size and, more directly, channel density modulate the detection threshold and resolution of cellular mechanosensing.

Still, what fraction of all channels are actually activated during a given mechanical stimulus still remains unclear. Mechanotransduction differs from, for example, voltage-sensing, where stimulus intensity is spatially homogenous. In contrast, forces are generally not uniform across the entire cell (Young et al., 2022). To answer this question, we focused on TREK1 for which we designed a new stimulation protocol. We used an internal buffer with Rb^+^ as the dominant ion, which has been shown to left-shift the voltage dependence of TREK1 (Schewe et al., 2016). In this buffer we were able to saturate the conductance-voltage relationship of TREK1 at 160 mV and thus, after normalizing for driving force, calculate the number of channels present in the entire cell (**Figure 4G, Figure 4—figure supplement 1**). Across all cells we measured, the total number of channels ranged roughly between 1,000 and 2,000. In the same experiment we mechanically stimulated cells, as previously described, and determine the number of channels recruited by the mechanical stimulus relative to the voltage step. Surprisingly, despite an overall large-scale compression, which elicited robust mechanically activated currents (161±22 pA; n=12), only 6.9±1.2% (n=12) of all channels were activated by this mechanical stimulus (**Figure 4H**). Importantly, the value of 6.9±1.2% may still be an overestimate since the mechanically gated open state of TREK1 has a two-fold higher conductance as compared to the voltage-activated state, although this was in standard K^+^ buffer (Rietmeijer et al., 2021).

In addition, we hypothesized that the cytoskeleton, which has previously been implicated in mechanoprotection of FGICs, but also in transmitting forces, contributes towards setting cellular detection limits (Cox et al., 2016; Gottlieb et al., 2012; Jia et al., 2016; Lotshaw, 2007; Shi et al., 2018; Verkest et al., 2022; Wang et al., 2022). We therefore decided to disrupt cytoskeletal architecture and measure which of these two opposing effects dominates. Specifically, we treated cells for 1 hour with Cytochalasin D (10 µM), which competes with barbed-end actin-binding proteins to affect both polymerization and depolymerization, destabilizing the overall actin architecture (Flanagan and Lin, 1980; MacLean-Fletcher and Pollard, 1980). Any reduction in *W*_*threshold*_ or *W*_*resolution*_ would point to a net mechanoprotective role of the actin cytoskeleton, whereas an increase would indicate mechanotransmission instead dominates. First, in separate AFM experiments we established that cytochalasin D treatment effectively reduces the elastic modulus (cells become softer): CytoD: 70.3±5.7 Pa (n=12), Untreated:147.8±20.8 Pa (n=12), p=0.005 (**Figure 5A-B**). We next applied our initial AFM-electrophysiology stimulation protocol to determine *W*_*threshold*_ and *W*_*resolution*_ on both Cytochalasin D treated cells and day-matched vehicle controls (**Figure 5C**). Cytochalasin D treatment led to a substantial decrease in the detection threshold (CytoD:13.7±1.3 fJ, n=16; DMSO: 34.9±6.0 fJ, n=14; p=0.0049) in cells expressing Piezo2 (**Figure 5D**). Similarly, work resolution decreased >4-fold as compared to controls (CytoD:0.08±0.02 fJ, n=16; DMSO: 0.38±0.09 fJ, n=14; p=0.0073) (**Figure 5E**).

**Figure 5.**
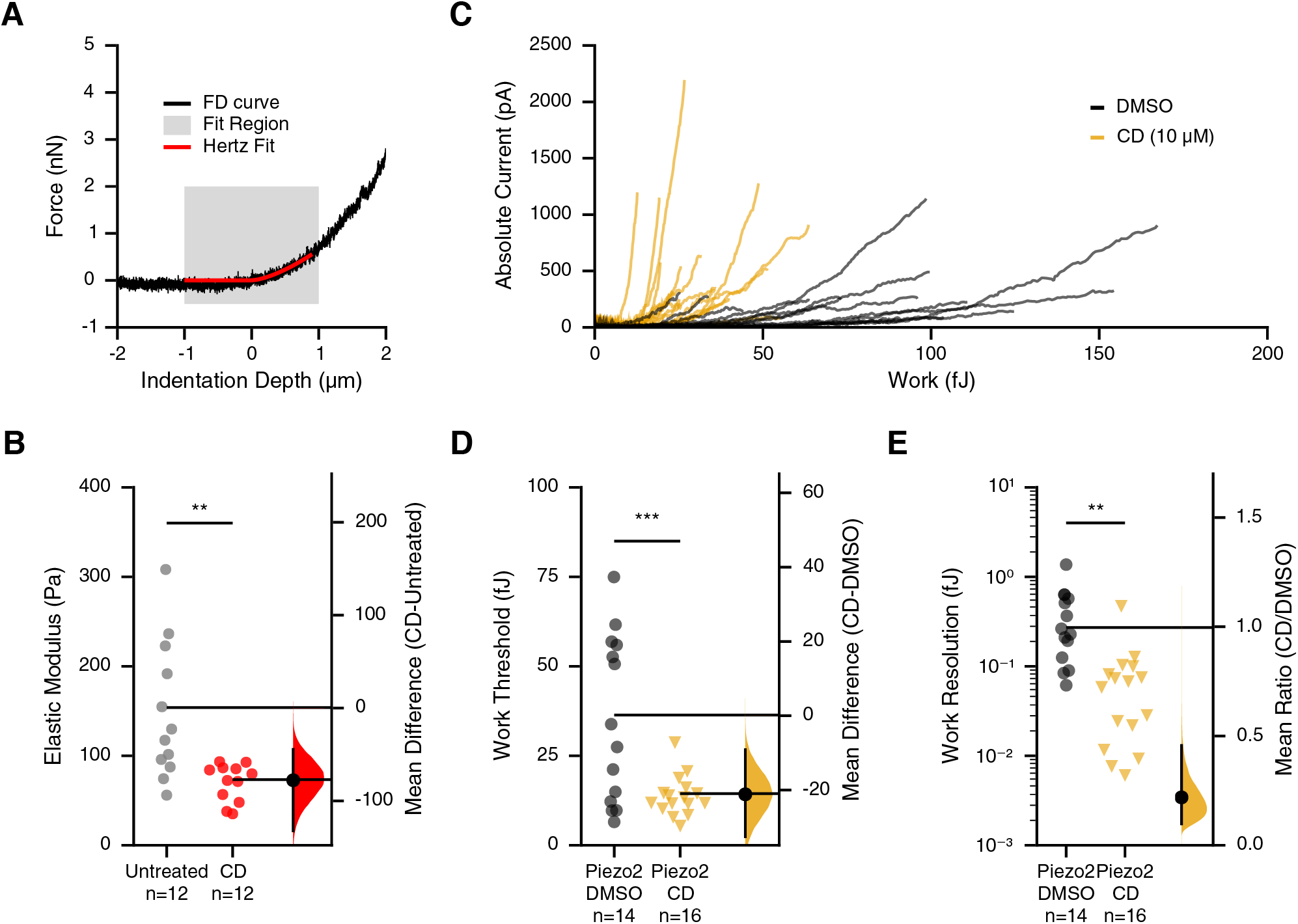
Piezo2 is sensitive to actin cortical disruption by cytochalasin D. **(A)** Representative force-distance curve (black) of a single cell with an overlaid Hertz model fit (red) to the first 1 µm of indentation. **(B)** Elastic modulus of Hertz model fits to individual HEK293T cells treated with 10 µm Cytochalasin D (red) or untreated control (gray). Group comparisons were performed using Welch’s T-test (p-value of untreated-CD: 0.005). **(C)** Raw work-current relationships for cells expressing Piezo2 and treated with 10 μM Cytochalasin D (orange) or DMSO control (black). Work threshold **(D)** and work resolution **(E)** values for cells expressing Piezo2 10 μM Cytochalasin D (triangles) or DMSO (circles). N indicates the number of replicates (individual cells). Group comparisons were performed using Welch’s T-test (p-values for work threshold of Piezo2+DMSO-Piezo2+CD:0.005 and for work resolution of Piezo2+DMSO-Piezo2+CD:0.007). The right side shows the bias-corrected and accelerated bootstrap distribution of mean differences **(D)** or ratios **(E)** between treated and control cells. Individual cells are shown as points, means are shown as horizontal lines, and vertical line corresponds to the 95% confidence interval of the bootstrap distribution.

Surprisingly, this effect was specific to Piezo2, as we saw no significant changes in *W*_*threshold*_ or *W*_*resolution*_ for cells expressing TREK1, TRAAK, or Piezo1 **(Figure 5—figure supplement 1)**, which suggests that Piezo2 is particularly sensitive to actin manipulation and that actin plays a predominantly mechanoprotective role. Taken together, we found that channel density is a general determinant of cellular sensitivity to mechanical forces, and that the actin cytoskeleton is a very effective modulator of cells expressing Piezo2.

### The speed of cellular mechanotransduction

It is known that Piezo and K2P channels open and conduct ions within milliseconds upon exposure to a mechanical stimulus (Berrier et al., 2013; Coste et al., 2010; Maingret et al., 1999). However, the precise timing of their response with respect to applied mechanical forces has not been quantified. In a subset of our collected recordings, we noticed a time delay (Δt) between the maximal applied force (peak force), which indicates the time point at which the piezoscanner ceases to move and compress the cell, and the resulting maximal transduction response (peak current) **(Figure 6A)**. Indeed, careful analysis revealed that cells expressing Piezo1 and TREK1 exhibited a substantial response delay of 8.2±2.2 ms (n=39) and 15.6±2.2 ms (n=60), respectively. Conversely, cells expressing Piezo2 and TRAAK responded within 1.5±0.5 ms (n=47) and 8.5±2.6 ms (n=38), respectively (**Figure 6B-C**). The rapid response of Piezo2 highlights that cells have, in principle, the ability to transduce forces with very fast (∼1 ms) speed. Further, for Piezo1 (and to a lesser extent Piezo2) the magnitude of the delay correlated with the intensity of the stimulus as measured by peak force amplitude (**Figure 6D-E**). This was not the case however for either TREK1 or TRAAK (data not shown). Importantly, we discovered that classical cell-indentation (poke) experiments on cells expressing Piezo1 or Piezo2 also revealed a response delay, after we aligned current responses to the end of the stimulus ramp (**Figure 6—figure supplement 1**), giving us confidence that this phenomenon was not an artefact of our AFM instrument.

**Figure 6.**
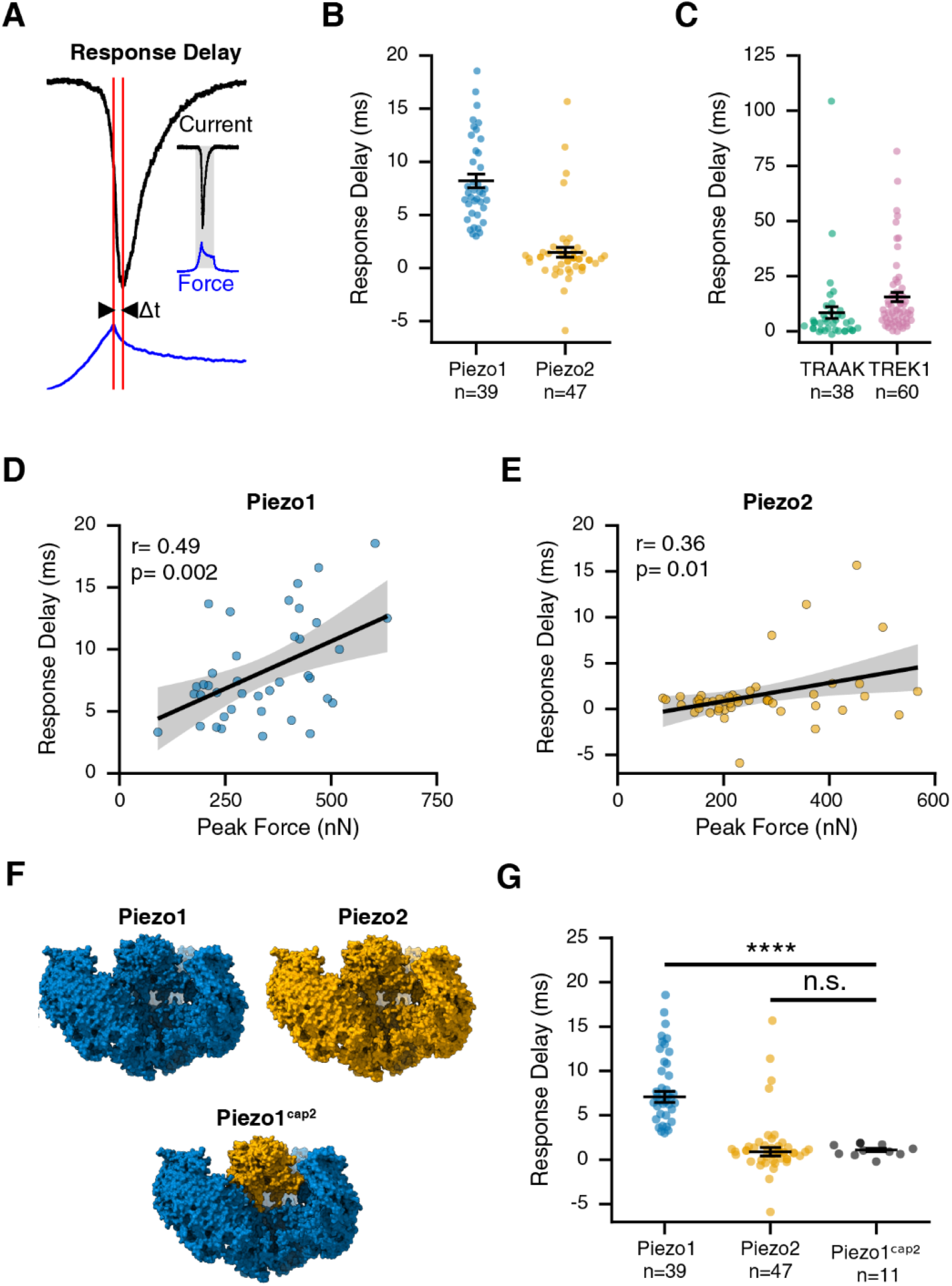
The speed of cellular mechanotransduction. **(A)** Calculation of response delay: representative traces of cantilever force (blue) and current (black) with red lines indicating the time difference (Δt) between peak force and peak current. Response delay values for cells overexpressing Piezo1 (blue) and Piezo2 (orange) **(B)**, and TRAAK (green) and TREK1 (pink) **(C). (D and E)** Response delay values as a function of peak stimulus force for Piezo1 **(D)** and Piezo2 **(E)**. Black lines are **l**inear fits and gray bands 95% confidence intervals. Correlation coefficients and p-values are indicated on the top left. **(F)** Schematic of chimeric Piezo construct (Piezo1^cap2^). **(G)** Response delay values for cells overexpressing Piezo1 (blue), Piezo2 (orange), and chimera Piezo^cap2^. Data for wild-type constructs are the same as in B. Group-wise comparisons were performed using Welch’s ANOVA followed by a post-hoc Games-Howell test (adjusted p-values for Piezo1-Piezo1^cap2^:1.17e-12 Piezo2-Piezo1^cap2^:0.728); n indicates the number of replicates (individual cells).

We hypothesized that both channel-intrinsic and cellular properties may account for this delay. We therefore first repeated our experiments with cells expressing a chimeric channel Piezo1^cap2^, for which previous work from our lab demonstrated that the cap domain of Piezo2 is sufficient to confer the fast inactivation kinetics of Piezo2 onto the normally slowly-inactivating Piezo1 (**Figure 6F**) (J. Wu et al., 2017). Indeed, cells expressing Piezo1^cap2^ responded within ∼1 ms (1.1±0.2 ms, n=11), which is statistically identical to Piezo2 (**Figure 6G**). Again, classical cell-indentation (poke) experiments produced qualitatively identical results (**Figure 6—figure supplement 1**). These results show that channel intrinsic properties, and in the specific case of Piezos, the cap domain, can determine the speed of cellular mechanotransduction.

Second, we reasoned that the slow diffusion of membrane tension may activate channels near the stimulation site, but with a time delay. We therefore turned to an *in silico* approach, where we simulated the gating of thousands of individual, spatially randomly distributed (∼100 channels/µm^2^), Piezo1 channels with a previously validated four-state Markov model (Lewis and Grandl, 2021) (Lewis and Grandl, 2015; Lewis et al., 2017). We then challenged channels with a spatially confined (2 µm radius) step in membrane tension, which diffused in two dimensions with a speed of either D = 0.024 µm^2^/s, which had been determined experimentally for HeLa cells, or 10 and 100-fold faster speeds (D = 0.24 µm^2^/s and 2.4 µm^2^/s, respectively), which is still one order of magnitude below the fastest diffusion values measured in neuronal axons (D = 20 µm^2^/s) (see Methods; **Figure 7A-C**) (Shi et al., 2022, 2018). Independent repetitions of this simulation with a diffusion constant of D = 2.4 µm^2^/s consistently showed a time delay in peak current amplitudes (dt = 6.2±0.6 ms) that approximated our experimental findings for Piezo1, while simulations with slower diffusion constants (D = 0 µm^2^/s, D = 0.024 µm^2^/s, and D = 0.24 µm^2^/s) produced time delays that were less pronounced or entirely absent **(Figure 7E)**. Overall, we conclude that the time delay we observed experimentally is indeed consistent with being an emergent property of membrane tension diffusion and further that the magnitude of this time delay is determined by the identity of the FGIC and the specific diffusion rate of the cell membrane.

**Figure 7.**
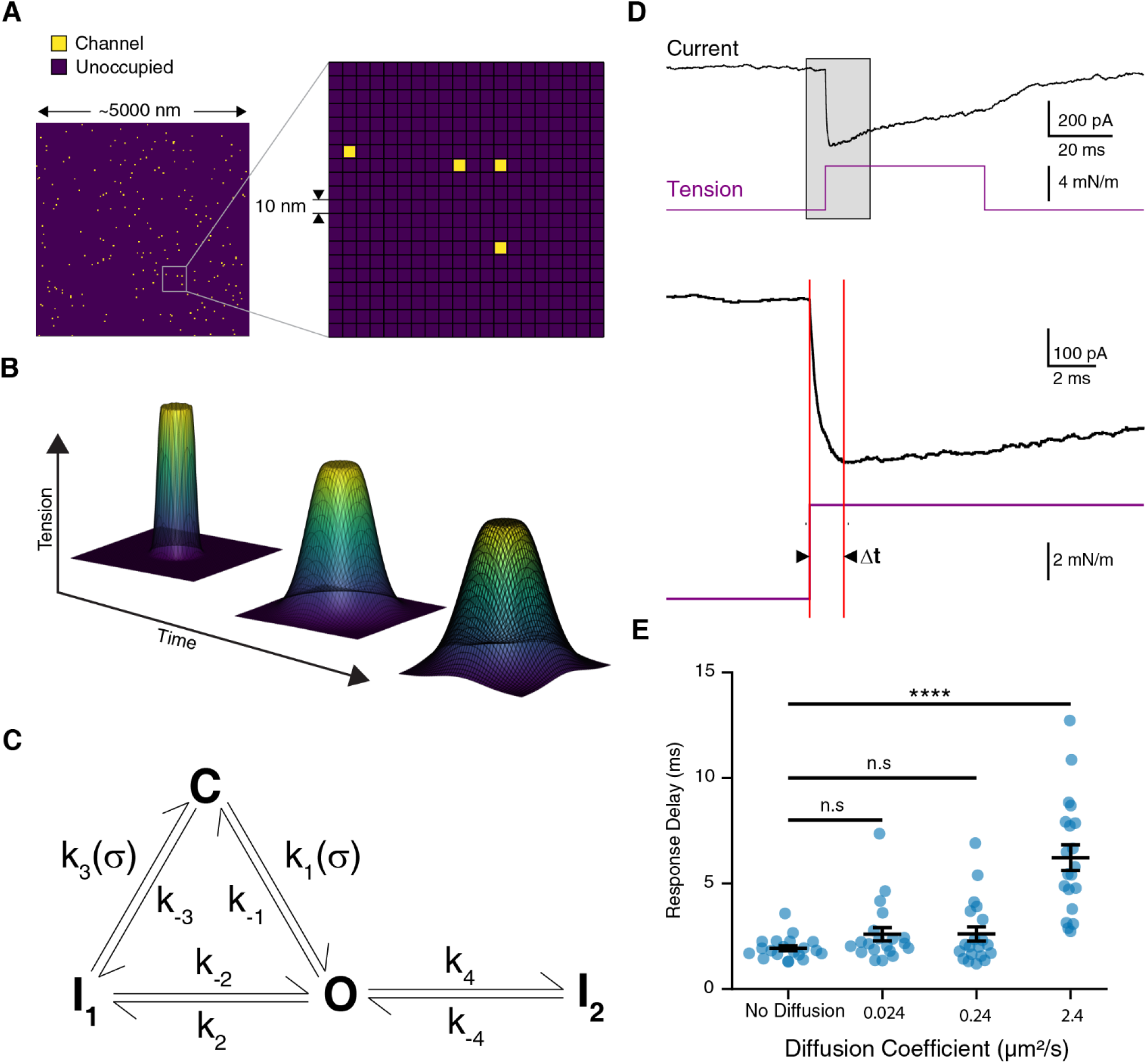
A model for simulating Piezo1 channel activation by membrane tension. **(A)** A representative 25 µm^2^ grid with randomly distributed channels (left). A 400 nm^2^ inset of the grid. Each unit square is 10 nm^2^ **(B)** Representative surface maps of tension diffusion as a function of time. Tension was clamped in a circular region of the grid and diffused outward. **(C)** Schematic of the kinetic rate model used in the Markov simulations. **(D)** Representative simulated Piezo1 current in response to a tension step protocol zoomed in to show the response delay. Red lines indicate the time of stimulus onset and peak of the current response. The full trace is shown in the inset. **(E)** Simulated response delay as a function of membrane tension. Twenty individual simulations were performed for each diffusion coefficient and are shown as blue points. Group-wise comparisons were performed using Welch’s ANOVA followed by a post-hoc Games-Howell test (adjusted p-values for No Diffusion-0.024 µm^2^/s: 0.226, No Diffusion-0.24 µm^2^/s:0.258, No Diffusion-2.4 µm^2^/s: <0.0001).

## Discussion

In this study, we aimed to explore the capabilities and limitations of cells as transducers of mechanical energy. To this end, we developed an instrument that combines AFM with whole-cell patch-clamp electrophysiology. This approach comes with caveats and potential sources for errors that need to be considered: First, we estimate an uncertainty in cantilever calibration of ≲ 20%, which is consistent with other studies (Song et al., 2015). Second, nonlinearity in the photodetector response may result in an underestimate of the most extreme forces of up to 29±3% and consequently also the calculated energies. This error has only a minimal effect on work threshold values, since these are determined within the linear range of the photodetector but may result in an overestimate of work resolution values. Third, the precise contact area between the cantilever and cell is unknown, which introduces an uncertainty in the spatial distribution of the stimulus as well as variability between measurements. Additionally, the time course of the stimulus (150 ms) is slow relative to the inactivation rates of the channels we studied. As a result, competing activation and inactivation lead to an underestimate of peak channel activity. Finally, by forming a whole-cell configuration the cell is no longer a hydrostatically closed system, and mechanical load may dissipate through the patch pipette, which again may lead to an overestimate of the work performed on the cell. Indeed, this may contribute to the force relaxation we observed during the holding phase of the cantilever. Still, the specific forces we measured at the transduction threshold for Piezo1 (228.2±16.4 nN; n=39) are comparable to those found using calcium imaging (185 nN), where the cell membrane remains intact, giving us confidence in these measuerments (Gaub and Müller, 2017). In any case, the detection threshold and work resolution we obtain for cells expressing different ion channels are clearly distinct, arguing that our instrument has sufficient precision.

In addition to experimental uncertainties, our experimental preparation differs from a physiological environment in several aspects: First, we seeded cells at low density and onto glass coverslips, which is necessary to both mechanically and electrically isolate cell properties, but removes contributions from neighboring cells present in tissue, and may alter the mechanical properties of the cell itself. For example, some evidence suggests that cell elasticity may vary with the stiffness of the underlying substrate, although a more recent study challenges this finding (Rheinlaender et al., 2020; Tee et al., 2011). In any case, by using glass, which has a stiffness that is much higher than that of the cantilever, substrate deformation and the related energy loss is minimized and therefore nearly all mechanical work is performed on the cell.

Second, we overexpress the channel of interest, which does not reflect native expression levels, but is necessary to overcome potential contributions of endogenous FGICs, and allowed us to clearly determine the role that channel identity plays in setting the physical limits (Dubin et al., 2017). Third, the flat cantilever of our AFM instrument is not a natural stimulus, but the resulting large-scale compression of the cell is arguably similar to the stimulation some cells-types experience *in-vivo*: For example, chondrocytes, which express both Piezo1 and Piezo and form cartilage of joints, and osteoblasts, which express Piezo1 and synthesize bones, are globally compressed by body weight (Hendrickx et al., 2021; Lee et al., 2014; Sun et al., 2019). Similarly, Merkel cells, which express Piezo2, are compressed by indentation of the skin (Maksimovic et al., 2014). Therefore, our approach is not only biophysically useful, but to some extent also physiologically relevant. Taken together, the approach and preparation we use are not intended to match physiological settings but are ideal for exploring the limits of cell-autonomous rapid force-sensing.

Biological sensory systems have evolved exquisite sensitivity to their relevant stimuli, while operating under constant noise present in the environment, which ideally enables reliable detection and discrimination of relevant signals. At the lower limit all physical systems are subject to thermal noise (k_b_·T = 4.11·10^−21^ J at room temperature) and FGICs themselves are estimated to have gating energies very near this thermal limit (Brohawn et al., 2014a; Cox et al., 2016; Lewis and Grandl, 2015; Lin et al., 2019; Maksaev et al., 2011). We were surprised to find that even in an overexpression system the lowest detection threshold and resolution we observed were 68.2±6.3 fJ and 1.0±0.2 fJ, respectively. For comparison, our own visual system can detect single photons (∼4·10^−19^ J), although only in low-noise conditions (dark) (Rieke and Baylor, 1998). The relatively high energy required for mechanical sensing may therefore be a consequence of a high magnitude of mechanical noise present in cells, for example caused by intrinsic forces generated by myosin motors (Ellefsen et al., 2019). In line with this interpretation is our result that only a small fraction (6.9±1.2%) of TREK1 channels is activated by our large-scale cell compression and that current responses never saturated. For Piezo channels this fraction may be higher, because despite a smaller unitary conductance (∼20-30 pS for Piezos vs ∼90 pS for TREK1) and comparable driving force (∼80 mV for Piezos vs ∼95 mV for TREK1), and faster inactivation, the elicited peak current amplitudes were higher. Regardless, our measurements clearly show that cells are inherently inefficient sensors of mechanical energies.

Three additional and very surprising findings emerged from our study: First, we found that cells can have pronounced (∼10 ms) delays in their transduction response. Our chimeric approach shows that response delay is a channel intrinsic property and that cells can inherently respond rapidly (<1ms). In addition, it was intriguing that this delay is almost absent in cells expressing Piezo2, which are present in DRG neurons, Merkel cells, and hair cells of the inner ear (Ranade et al., 2014b; Woo et al., 2015, 2014; Z. Wu et al., 2017). It is therefore interesting to speculate that Piezo2 may have specialized to respond rapidly to fulfill its role in rapid sensory transduction. Second, our simulations show that a time delay can emerge naturally from the slow diffusion of tension. Of course, the actual spatial distribution of the stimulus and the speed of membrane diffusion that underlie our experiments are likely different from the tension clamp we simulated, which may explain the less pronounced delays *in silico*. Moreover, differing tension sensitivities and inactivation kinetics may explain why we found that Piezo2, TREK1, and TRAAK have response delays that are distinct from Piezo1. For example, increased tension sensitivity or slower inactivation kinetics may easily explain how channels can be activated with high probability far away from the stimulation site, altogether resulting in longer response delays. Importantly, recent work has shown that certain subcellular domains may be specialized to diffuse tension much faster, allowing long-range mechanical coupling. Interestingly, neuronal axons are among these regions which may have important implications for the transduction of mechanical touch (Gomis Perez et al., 2022; Shi et al., 2022). We propose that in addition to long range mechanical coupling, tuning the rate of tension diffusion has important implications for the timing of the mechanotransduction response. This property may be particularly important for sensory systems, where the cellular response needs to reliably reflect the temporal characteristics of the underlying stimulus.

Third, we found that cells expressing TRAAK and TREK1 function as proportional transducers of mechanical energy, whereas cells expressing Piezos showed a highly nonlinear step-like response. Qualitatively, this behavior is reminiscent of the respective pressure-clamp (stretch) induced responses for TRAAK, TREK1, and Piezo1 (Piezo2 is not efficiently activated by membrane stretch), which have shallower slopes in their pressure response curves relative to Piezo1 (Brohawn, 2015; Brohawn et al., 2014b; Coste et al., 2010). On a cellular level this response property has important implications for how stimulus intensity is encoded. For example, a shallow, graded response is better suited to encode stimuli across a wider dynamic range, whereas a steep, switch-like response is better suited to encode the crossing of a particular threshold. Interestingly, Piezo1 showed a particularly steep work-current relationship, which raises the question how this widely expressed ion channel may endow different cell types to sense distinct mechanical stimulus intensities.

We also identified several factors that influence the response properties of cells. Our data suggest that cell size and channel density may be mechanisms to modulate the response properties of cells. The fact that we only observed a correlation between cell size and response properties in cells expressing Piezos, but not in K2P channels may be due to differences in dynamic range and variability in expression levels we are able to explore. Additionally, disrupting the cytoskeleton resulted in a stark decrease in both the detection threshold and the work resolution, specifically in cells expressing Piezo2. This result is consistent with the cytoskeleton playing a strong mechanoprotective role by limiting the transmission of mechanical energy to Piezo2 channels. It would be interesting to see, given the role of Piezo2 in sensory transduction, whether specialized sensory cells take advantage of this sensitivity to actin disruption, for instance by localizing Piezo2 to specialized signaling domains of low actin density.

Altogether, our novel approach and its results provide a quantitative framework for understanding the energetics of mechanosensing on the level of single cells. A natural extension of this work would be to next characterize primary mechanosensory cells. From there, we may learn the magnitudes of mechanical energies a particular cell-type is tuned to sense, how they compare to the parameter space we found in our heterologous system, and finally what mechanisms these cells may have to extend or constrain their properties as cellular mechanosensors.

## Acknowledgements

This study was supported by NIH 5R01NS110552 (M.Y. and J.G), The Ruth K. Broad Biomedical Research Foundation (M.Y.) and the Duke Institute for Brain Sciences (DIBS). We thank Marie Cronin for thoughtful comments on the study, and Tejank Shah for technical assistance.

## Materials and Methods

### Cell Culture

HEK293T cells (ATCC #11268 authenticated and tested mycoplasma free, Manassas, VA) were cultured at 37 °C and 5% CO_2_ in DMEM-HG (Life Technologies, Carlsbad, CA) supplemented with 10% FBS (Clontech, Mountain View, CA), 50 U/mL penicillin, and 50 mg/mL streptomycin (Life Technologies). Cells were reseeded into 6-well plates and transfected (mouse Piezo1-pIRES-EGFP (Coste et al., 2012), mouse TREK1-pIRES2-EGFP, and mouse pCEH-TRAAK-GFP, YFP alone: 3 µg; mouse Piezo2 (Lewis et al., 2017): 2.5 µg + 0.5 µg YFP) with Fugene 6 (Promega, Madison, WI) 48 hours prior to recording at a transfection ratio of 10 µL Fugene6:3 µg total DNA. Glass coverslips (1.5 mm; Warner Instruments, Hamden, CT) were coated with 0.1 mg/mL Poly-L-Lysine at room temperature for 30 minutes followed by 1 µg/mL Laminin at 37 °C for 45 minutes to 1 hour. Cells were trypsinized and transferred onto coated coverslips16-24 hours before recording. Cell cultures were limited to 20 passages.

Cells treated with Cytochalasin D (10 μM solubilized in DMSO; both Sigma, Burlington, MA) for 1 hour prior to and throughout the recording session. Control cells were day-matched and treated with an equal volume of DMSO over the same time course.

### AFM Instrument

A Digital Instruments (Bruker, Billerica, MA) Bioscope was modified for simultaneous use with electrophysiology: A custom-designed aluminum stage was manufactured (Protolabs, Maple Plain, MN) to mount the Bioscope to a Nikon Ti-E inverted microscope. The position limiters on the stock Bioscope head were removed and a custom stopper was designed and 3D-printed to increase the lower-limit of the Bioscope head position. The stock fluid-cell probe holder was BK-7 glass with anti-reflective coating and custom cut (Mindrum Precision, Rancho Cucamonga, CA) to allow access to a patch-pipette near the cantilever tip. The pipette manipulator was set to a 15º angle, and the stage (Nikon, Tokyo, Japan) was additionally modified to increase the lower limit of the manipulator position. A large bath was 3D printed to allow positioning of both the probe-holder and a patch-pipette over the coverslip. The photodetector signal was passed from the Bioscope through a signal access module (Veeco, Plainview, NY) to an EPC10 amplifier (HEKA Elektronik). The signal access module also allowed external control of the piezoscanner. For this, a digital trigger signal was passed from Patchmaster (HEKA Elektronik) to custom software written in Labview 2016 (National Instruments, Austin, TX) generating the stimulus waveform, which was then passed to a high-voltage amplifier (EPA-104; Piezo.com, Woburn, MA) and subsequently via the signal access module to the Nanoscope3a controller (Bruker) and the piezoscanner z-drive. The correspondence between the scanner position and the applied command voltage was determined by the scanner calibration in the Nanoscope software (v5.33) after manufacturer calibration.

### AFM Calibration

CSC37, CSC38, or NSC35 tipless, chromium and gold-coated cantilevers (Mikromasch, Tallinn, Estonia) were mounted to a custom probe holder and attached to the Bioscope head. Prior to experiments, for each individual cantilever the stiffness was determined using the method of Sader (Chon et al., 2000; Sader, 1998; Sader et al., 1999). Briefly, the dimensions of the cantilever were measured optically (Nikon Ti-E). The quality factor (Q), amplitude (A), and resonant frequency (f_0_) were determined by fitting the equation for a simple harmonic oscillator to power spectra of the free cantilever vertical displacement signal (P) in air averaged across 512 spectra using Welch’s method (Welch, 1967) and sampled at > 5-fold the expected resonance frequency using a National Instruments Data Acquisition Board (NI-USB 6361).

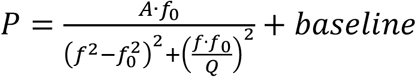

The resonant frequency (f_0_), quality factor (Q), cantilever length (L), and width (w) were used to determine the stiffness of the cantilever (k) using the following relationship:

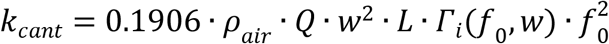

where ρ_air_ is the density of air at 25 °C (1.18 kg/m^3^) and Γ_i_(f_0_, w) is the imaginary component of the hydrodynamic function. The photodetector sensitivity was calibrated by pressing the mounted cantilever onto bare glass prior to stimulating a nearby cell and using the inverse optical lever sensitivity (invOLS) method (Gavara, 2017): Briefly, due to the high stiffness of the coverslip relative to the cantilever the slope of the relationship between the vertical difference signal of the photodetector and the piezoscanner position is defined to be 1. Thus, the inverse of the measured slope equals the sensitivity. We calculated the average sensitivity from 10 individual measurements. All sensitivity calibrations were performed in standard recording buffers prior to measuring each cell. We estimated the uncertainty in our measurements on a subset of 20 cells based on the uncertainty in the fit parameters and the variance in the sensitivity calibration.

### AFM Stimulation of HEK293T Cells

All stimuli were applied at 5 s intervals. Prior to initial contact, a coarse step-motor moved the cantilever into contact with the cell in 1-5 µm steps. Contact was determined by a sudden increase in the photodetector signal. Following contact, the step-motor moved the cantilever towards the cell in 0.2-1 µm increments until mechanotransduction currents were observed. Stimuli were applied at a rate of 40 μm/s for a total distance of ∼6 μm. The speed was chosen based on initial validation experiments in HEK293T cells overexpressing Piezo1 as it was the slowest speed that was able to elicit robust mechanotransduction currents of the speeds tested (10 μm/s, 20 μm/s, 40 μm/s, 80 μm/s, 160 μm/s).

### Electrophysiology

All whole-cell recordings were performed at room temperature using an EPC10 amplifier and Patchmaster software (HEKA Elektronik). Data were sampled at 25 kHz and filtered at 2.9 kHz using an 8-pole Bessel filter. Series resistance was compensated 20-65%. Thin-walled borosilicate glass pipettes (1.5 mm OD, 1.17 mm ID; Sutter Instrument Company, Novato, CA) were pulled and wrapped in parafilm to reduce pipette capacitance. The final pipette resistance was 2.0-6.0 MΩ when filled with pipette solution (in mM: 133 CsCl, 0.5 EGTA, 10 HEPES, 1 MgCl_2_, 4 MgATP, 0.4 Na_2_GTP for Piezo1, Piezo2, and YFP or 133 KCl, 0.5 EGTA, 10 HEPES, 1 MgCl_2_, 4 MgATP, and 0.4 Na_2_GTP for TRAAK and TREK1). The bath solution for experiments involving Piezo1, Piezo2, and YFP contained in mM: 150 NaCl, 1 MgCl_2_, 2.5 CaCl_2_, 10 HEPES, 10 Glucose, and 3 KCl), and the bath solution for experiments involving TRAAK or TREK1 contained in mM: 140 NaCl, 3 MgCl_2_, 1 CaCl_2_, 10 HEPES, 3 KCl, 10 Glucose, and 10 TEA-Cl. For TREK1 experiments using RbCl-based solutions the bath solution consisted of in mM: 140 NaCl, 3 MgCl_2_, 1 CaCl_2_, 10 HEPES, 30 KCl, 10 Glucose, and 10 TEA-Cl. The pipette solution for these experiments consisted of in mM: 133 RbCl, 1 MgCl_2_, 10 HEPES, 0.5 EGTA, 4 MgATP, and 0.4 Na_2_GTP. For bath solutions pH was adjusted to 7.4 and for pipette solutions 7.2 using the hydroxide of the dominant cationic species. Osmolality was adjusted to ∼310 for bath solutions and ∼290 for pipette solutions with sucrose when necessary. The internal solution was allowed three minutes to dialyze prior to recording to promote GTP-mediated run-up of Piezo currents (Jia et al., 2013). Recording sessions for individual coverslips were limited to one hour.

### Elasticity Measurements

Cell elasticity measurements were performed with gold and chromium coated cantilevers with ∼10 µm borosilicate colloids probes (CP-qp-CONT-BSG: NanoandMore USA, Watsonville, CA). The exact probe diameter was measured using an FEI Apreo Scanning Electron Microscope (SEM) and averaged from the major and minor diameter of an overlayed ellipse. The cantilever was calibrated as described above with the exception that only a single invOLS measurement was performed at the start of each measurement. The cantilever was displaced a total distance of ∼6 μm at a rate of 1 µm/s before being immediately retracted at the same speed. A linear fit was performed on the first 20% of the data, prior to contact, and subtracted from the trace. The contact point was determined, based on visual inspection, as the inflection point of the force-distance curve. A Hertzian contact model was fit to the first 1 μm of the force-distance curve following the contact point and used to estimate the elastic modulus (E) as follows:

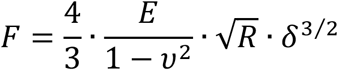

where R is the radius of the probe determined by SEM imaging, *υ* is the Poisson ratio taken to be 0.5, and *δ* is the indentation depth.

### Analysis

All data were analyzed using custom analyses written in R, Python, and Julia (Github: GrandlLab). Cells with leak currents >300 pA at -80 mV or series resistances >15 MΩ were excluded from the analysis. Cells were determined to be responsive to mechanical stimuli if they displayed stimulus-locked whole-cell currents of *≧* 50 pA. Baseline currents and photodetector voltage in the absence of mechanical stimulation were subtracted offline. For TRAAK and TREK1 the mechanocurrent was isolated from the voltage-dependent current by subtracting the current amplitude at 0 mV. The compression force (F) was calculated from the baseline subtracted vertical difference signal of the photodetector (V_PD_) using the following equation, where S denotes the sensitivity and k_cant_ denotes the cantilever stiffness:

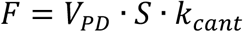

The distance traveled by the cantilever was corrected to account for cantilever deflection using the following equation:

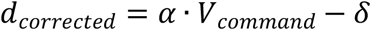

where α is the calibration factor for the piezoscanner, V_command_ is the command voltage sent to the photodetector, and *δ* is the deflection of the cantilever.

The work was determined by calculating the cumulative integral of the calculated force as a function of the distance traveled by the cantilever.

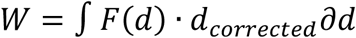

For analyses of the approach phase of each stimulus, only data during the initial ramp of the cantilever was considered.

Transduction characteristics were determined as follows:

1. Power coefficients for individual work-current curves were determined by removing data prior to the work threshold, log transforming, and performing a linear fit.
2. Work threshold (*W*_threshold_) values were determined as the maximum work, at which the current remains below 2 standard deviations above the baseline current:

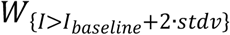

where the standard deviation was calculated over a 100 ms window preceding stimulus onset. Average values for 2 · *Standarddeviation* were 4.0±0.5 pA for Piezo1, 4.4±0.4 pA for Piezo2, 7.5±0.7 pA for TRAAK, and 12.8±0.8 pA for TREK1. Current threshold values for TREK1 and TRAAK are likely slightly higher due to their higher open probability at 0 mV. Current traces were passed through an additional 6-pole Bessel filter with a cutoff frequency of 1 kHz offline prior to threshold determination.
3. Work resolution (W_resolution_) values are calculated from the inverse slope (dW/dI), by performing a linear least square fit to the range of data corresponding to the steepest region of the work-current relationship, and the following equation:

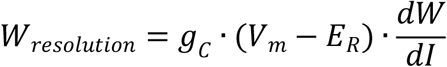

using previously reported single channel conductances (g_C_) (Piezo1: 29.1 pS, Piezo2: 23.7 pS; (Coste et al., 2015), TRAAK: 65.4 pS, TREK1: 88.5 pS; (Blin et al., 2016). The reversal potential (E_R_) was estimated to be zero for Piezo channels and the Nernst potential for potassium (−95 mV) in the solutions used for K2P channels.
4. Channel density was estimated using the mean current (I_v_) at 0 mV in the absence of a mechanical stimulus and the membrane capacitance (C_m_) using the following equation.

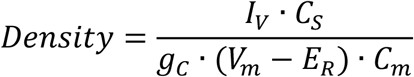

where g_c_ is the respective single-channel current at the applied voltage and C_S_ is the approximate capacitivity of biological membranes of 1 µF/cm^2^ (Gentet et al., 2000).
5. The total number (N) of TREK1 channels was calculated using the following equation:

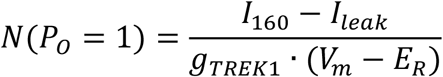

where I_160_ is the current at 160 mV, I_Leak_ current obtained from a *post hoc* linear fit to currents evoked by increasing negative steps from -80 mV to -90 mV to -80 to -120 mV, g_Trek1_ the unitary conductance for TREK1 in K^+^ buffers (Blin et al., 2016), and E_R_ the reversal potential in Rb^+^ buffer determined with a tail-current protocol **(Figure 4—figure supplement 1)**.
6. The number of channels activated by a mechanical stimulus (N_mech_) was calculated by additionally subtracting the voltage-activated current (I _v_) at 0 mV from the mechanically activated current (I_mech_) and using the following equation:

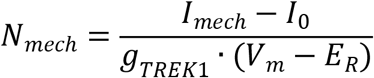
7. The response delay (Δt) in AFM experiments was calculated as the time difference between the peak of the mechanically activated current 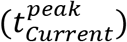 and the peak of the applied force 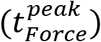:

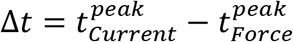
8. The response delay (Δt) in traditional poke experiments was calculated as the time difference the peak of the mechanically activated current 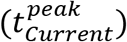 and the peak of the stimulus ramp 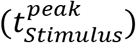:

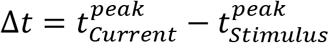

All values reported are bootstrapped means±sem unless otherwise noted. All statistical analyses were performed using Welch’s t-test for two group comparisons. For omnibus tests Welch’s ANOVA was performed followed by a post-hoc Games-Howell test. Significance thresholds correspond to: * p < 0.05, ** p < 0.005, *** p < 0.001. Bootstrap distributions and confidence intervals were determined using the bias-corrected and accelerated bootstrapping method except where SEM is reported in which case percentile bootstrap distributions were used. All bootstrapping was done with >=10,000 replicate.

### Markov model simulations

Channel gating was simulated with a custom-written script using Python available on GitHub (Github: Grandllab). Piezo1 channels were distributed randomly on a grid with 10 nm x 10 nm pixel-size. The grid size was adjusted for each simulation to minimize simulation time and to ensure tension at the edge never exceeded 5% of the maximal tension amplitude to minimize boundary effects. Pixel occupancy was approximately 1%, corresponding to the experimentally obtained channel density of Piezo1 overexpressed in Neuro2A cells of 100 channels/μm^2^ (Lewis and Grandl, 2021). Gating of each individual Piezo1 channel was simulated with a four-state Markov-model (**Figure 7C**) as previously described and adapted to be a function of membrane tension (σ) (Lewis and Grandl, 2015; Lewis et al., 2017; Young et al., 2022). Specifically, rate constants for wild-type Piezo1 (σ_50_ = 1.4 mN/m, slope factor b = 0.8 mN/m) were k_1_(σ) = 5.1·exp(σ/b) s^-1^, k_-1_ = 5·exp(σ_50_/b) s^-1^, k_2_ = 8.0 s^-1^, k_-2_ = 0.4 s^-1^, k_3_(σ) = 34.6·exp(-σ/b) s^-1^, k_4_ = 4.0 s^-1^, k_-4_ = 0.6 s^-1^, where σ is the applied tension, and k_-3_ = (k_1_·k_2_·k_3_)/(k_-1_·k_-2_) to achieve microscopic reversibility. Sampling intervals (dt) were adjusted such that dt << 1/k_1_(σ) at all times and ranged from 0.1 to 10 μs. Equilibrium occupancies, at σ=0, were determined analytically and used as starting values. Each simulation was allowed an additional 50 ms at σ = 0 to ensure complete equilibration, before the tension stimulus was applied. At *t* = 50 ms a circular tension stimulus with radius = 4 μm (200 pixels) was applied. Propagation of membrane tension was calculated with a 2D-Gaussian diffusion function (**Figure 7B**):

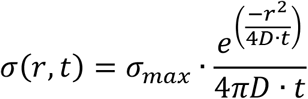

where σ_max_ is the maximal tension amplitude, D the diffusion coefficient, r is the distance of the channel from the edge of the stimulus, and t is the time after stimulus onset. Initially, 1 s simulations were performed to validate the baseline current, peak current, inactivation kinetics, plateau current matched expected values. Shorter simulations (90 ms) were produced to calculate response delay as a function of diffusion coefficient. Delay times were calculated by determining the time post-stimulus onset, at which the maximum number of opened channels occurred. Stimulations were repeated 10 times for each condition. All data are mean+sem.

## Data Availability

All code associated with cantilever calibration, photodetector calibration, data analysis, and simulation are available and accompanied by reproducible environments and documentation on the laboratory GitHub page (https://github.com/GrandlLab). Source data files have been provided with the numerical data for each figure. Raw data is hosted on Dryad and can be accessed via the following repository https://doi.org/10.5061/dryad.gtht76hq5.

---

**Figure 2—figure supplement 1.**
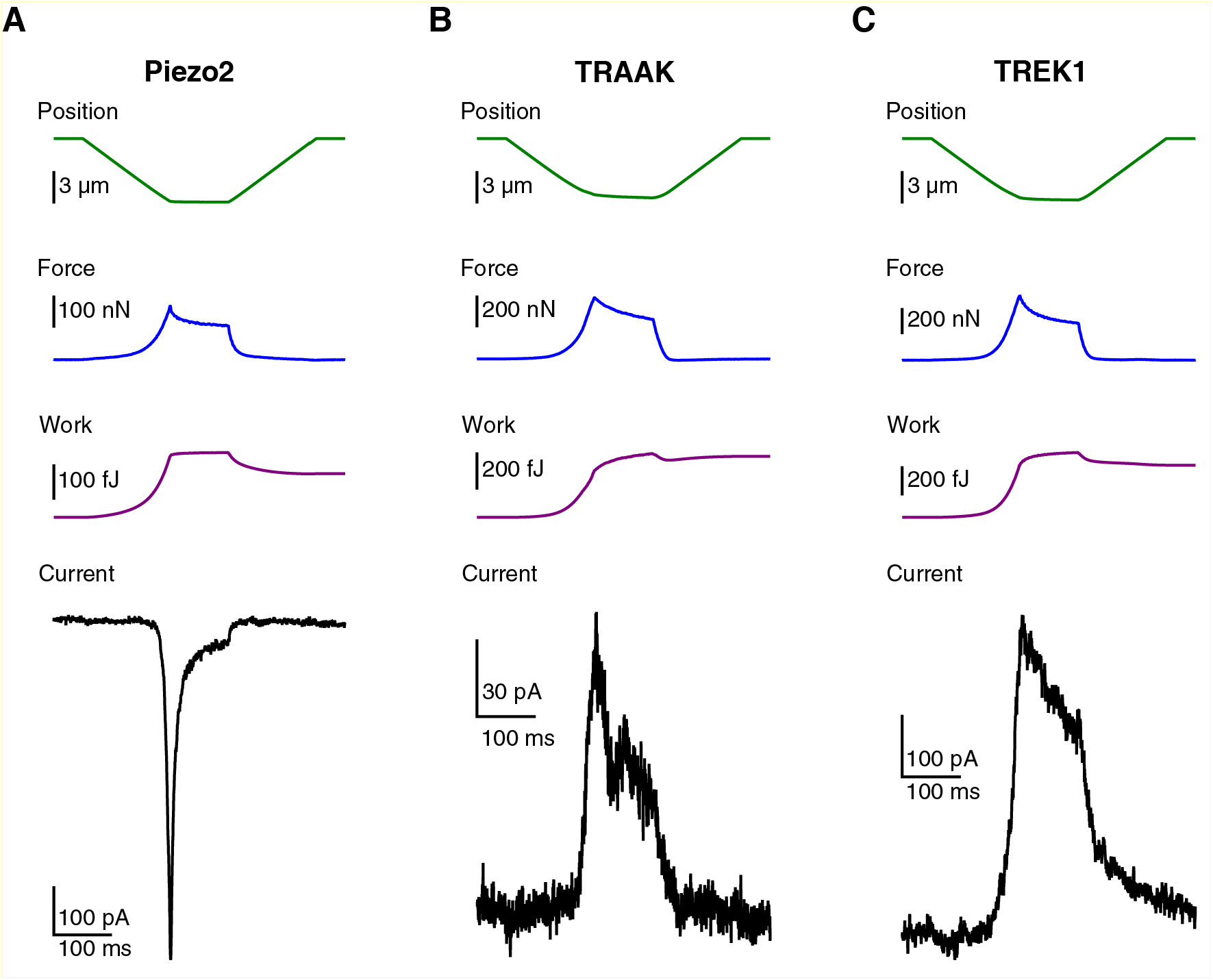
Representative Traces for Piezo2, TRAAK, and TREK1. **(A)** Representative trace for Piezo2 at -80 mV. **(B)** Representative trace for TRAAK at 0 mV. **(C)** Representative trace for TREK1 at 0 mV.

**Figure 4–figure supplement 1.**
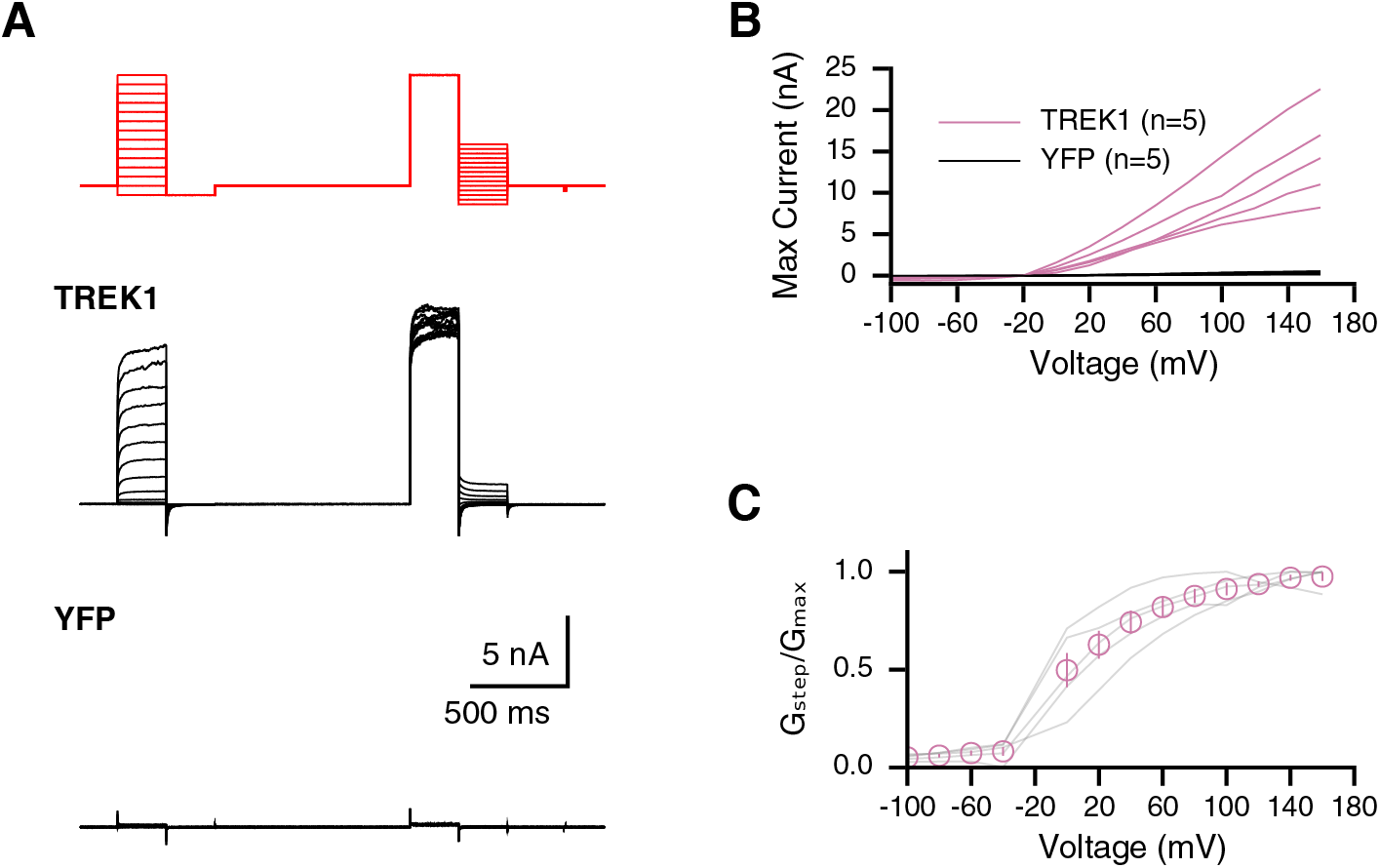
TREK1 voltage-dependence saturates in Rb^+^ internal solution. **(A)** Voltage-step and tail-current protocol and representative traces of cells overexpressing TREK1 or YFP. **(B)** Individual current-voltage relationships from cells expressing TREK1 (pink) or YFP (black). N indicates the number of replicates (individual cells). **(C)** Normalized average conductance-voltage relationship for cells shown in B.

**Figure 5—figure supplement 1.**
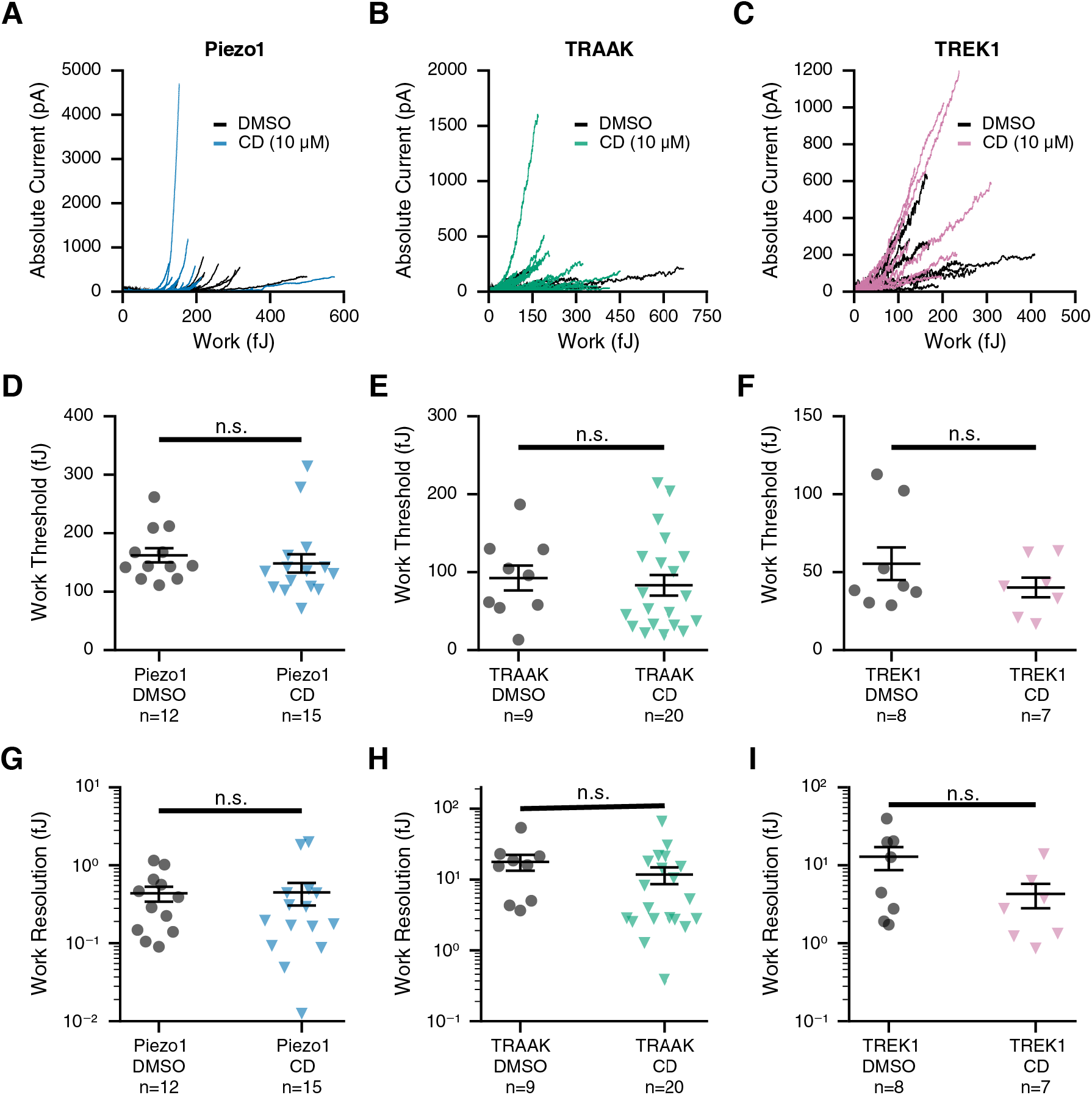
Piezo1, TRAAK, and TREK1 detection threshold and resolution are insensitive to Cytochalasin D treatment. Representative work-current relationships for Piezo1 **(A)**, TRAAK **(B)**, and TREK1 **(C)**. DMSO-treated control cells are shown in black, and cells treated with 10 µM Cytochalasin D are shown in blue, green, or pink respectively. Work threshold values for Piezo1 **(D)**, TRAAK **(E)**, and TREK1 **(F)**. Work resolution values for Piezo1 **(G)**, TRAAK **(H)**, and TREK1 **(I)**. Colors for D-I are as in A-C. For D-I, lines indicate the bootstrapped mean and SEM. N indicates the number of replicates (individual cells). Group comparisons in D-I were performed using Welch’s T-test.

**Figure 6–figure supplement 1.**
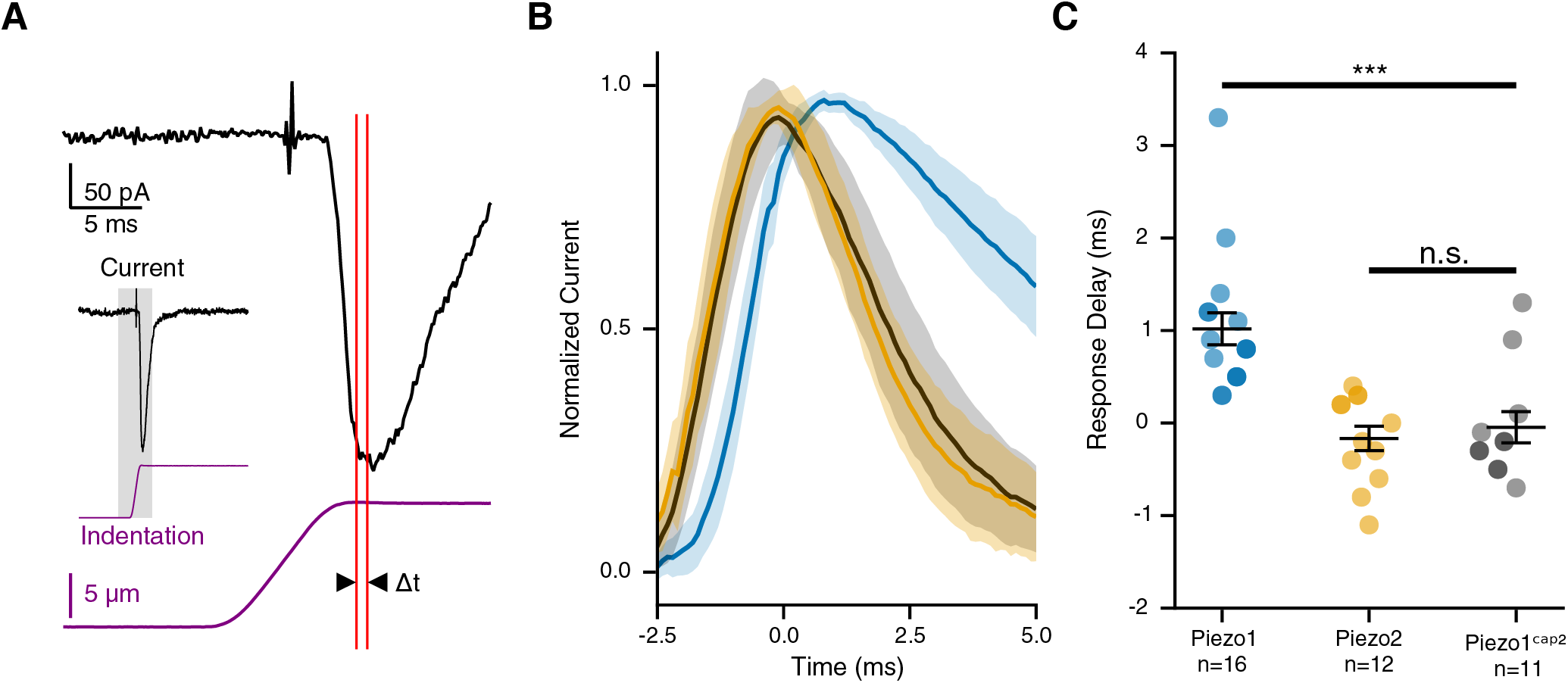
Cells expressing Piezo1 show a response delay in whole-cell poke stimulation. **(A)** Zoomed-in representative poke current trace (black) and stimulus (purple). Response delay (Δt) was determined based on the time difference between the peak of the current response and the end of the actuator displacement both shown in red). A zoomed-out view of the trace is shown in the inset with the region of interest highlighted in gray. **(B)** Average current traces upon whole-cell poke stimulation for cells overexpressing Piezo1 (blue), Piezo2 (orange), and chimera Piezo1^cap2^. Traces were normalized to their peak and aligned to the end of the end of the actuator displacement of the poke stimulation. Bands are 95% confidence intervals. **(C)** Response delay values, reflecting the time difference between the stop of the poke stimulation and the peak of the transduction current, for cells overexpressing Piezo1 (blue), Piezo2 (orange), and chimera Piezo1^cap2^. N indicates the number of replicates (individual cells). Group-wise comparisons were performed using Welch’s ANOVA followed by a post-hoc Games-Howell test (adjusted p-values for Piezo1-Piezo1^cap2^:0.001 and Piezo2-Piezo1^cap2^:0.861).

